# A Respiratory Syncytial Virus trailer sequence modulates viral replication and copy-back defective viral genome generation and propagation kinetics

**DOI:** 10.1101/2025.09.24.678299

**Authors:** Justin W. Brennan, Gaochan Wang, Sarah Connor, Xingjian Wang, Thomas J. Mariani, Yan Sun

**Author notes:** Correspondence, 601 Elmwood Avenue Rochester, NY 14642.

## Abstract

Copy-back defective viral genomes (cbDVGs) are key inducers of antiviral responses during negative-sense RNA virus infection. Once considered byproducts of *in vitro* viral replication, cbDVGs have since been detected in clinical specimens and implicated in affecting infection outcomes. The molecular mechanism of cbDVG generation remains unclear, thereby hindering our ability to manipulate cbDVG production during infection for therapeutic gain. Previous work showed that respiratory syncytial virus (RSV) cbDVG re-initiation sites cluster in trailer-end hotspots R1, R2, and R3, and that a poly-U mutation in R1 selectively reduced cbDVG formation at the mutated region. Here, we reported that a 10U mutation in R2 drastically reduced cbDVGs in this region in both minigenome and recombinant virus systems. Furthermore, during high-MOI passaging of the R2-10U virus, we observed delayed detection of cbDVGs with re-initiation sites in R1-R3 (trailer cbDVGs) compared to WT, while no differences in virus titers were observed. Interestingly, we observed the rapid emergence and accumulation of a viral variant bearing a 2-ribonucleotide deletion (R2-8U) within the R2-10U mutation sequence as early as P0. Compared to R2-10U, the R2-8U virus was stable, displayed faster generation and accumulation of trailer cbDVGs, restored cbDVGs with R2 re-initiation sites, and exhibited enhanced genomic replication. Overall, our data identify a sequence in the RSV trailer whose mutation critically modulates both viral replication and the generation/propagation of trailer cbDVGs. Our data also suggest that cbDVG generation, particularly near the trailer, may be an evolutionary tradeoff for more rapid virus genomic replication.

## Introduction

Defective viral genomes (DVGs) are truncated derivatives of their parental viruses generated during an aberrant round of viral genomic replication and are unable to replicate in the absence of a co-infecting, homologous, standard virus. DVGs have been observed during the infections of many RNA viruses (1–8), are well documented triggers of antiviral innate immune responses and certain DVGs interfere with the replication of their standard viruses by competing for critical viral proteins required for genomic replication or packaging into the virus particle (1, 3, 6, 9–19). Recent reports have detected DVGs in clinical samples from individuals infected with SARS-CoV-2, influenza A, dengue, measles and respiratory syncytial virus (RSV) (3, 14, 20–24). Importantly, the amount and kinetics of DVG production are associated with infection outcomes (23, 25). For example, a non-pathogenic influenza A virus, identified from mild human infections, accumulated high levels of DVGs *in vitro* and exhibited reduced pathogenicity in mice (25). Likewise, a longitudinal cohort study of adults experimentally infected with RSV showed that individuals with late or prolonged detection of DVGs had worse symptom severity scores and higher viral loads than individuals with early DVG detection, highlighting the critical role of DVG kinetics in influencing the morbidity of viral infection (23). Despite intensive studies on the functions of DVGs, the molecular basis of DVG generation is largely unknown.

Two major classes of DVGs, deletion- and copy-back (cbDVG) DVGs, are consistently observed during viral infections. While positive-sense RNA viruses and influenza viruses produce the deletion type of DVG (3, 24, 26–29), single-stranded negative-sense RNA viruses, like RSV, majorly produce the copy-back type of DVG (8, 21–23, 30, 31). cbDVGs are generated during an aberrant round of viral genomic replication when it is thought that the viral polymerase dissociates from the antigenomic template strand at a “break point” and reinitiates RNA synthesis at a “rejoin point” on the nascent strand and continues elongating through the 5’ end (32, 33). This leads to the generation of a theoretical panhandle RNA species containing reverse complementary termini and a unique break and rejoin junction sequence (34–36). We previously found that RSV cbDVGs with rejoin points located near and within the trailer complement sequence, spanning the last ∼200 nucleotides of the antigenome, are selectively enriched during high multiplicity of infection (MOI) passaging *in vitro* (8). In virus stocks enriched with these trailer cbDVGs, cbDVGs induce robust types I and III interferon (IFN) responses, reduce viral load, and alleviate lung pathology *in vivo* (13). Notably, trailer cbDVGs have also been identified in nasal samples from hospitalized pediatric patients with high viral loads (14). To investigate the viral factors that determine trailer cbDVG generation and accumulation during infection, we divided the antigenomic region clustered with cbDVG rejoin points into three hotspots: R1, R2 and R3. R1 spans the end of the L gene, including the L gene end signal, while R2 and R3 are located within the trailer complement sequence. Previously, we mutated a short sequence containing G/C ribonucleotides to a poly-U sequence within R1 and found that this mutation reduced the abundance of cbDVGs containing rejoin points within R1, suggesting that the composition of cbDVG populations can be genetically manipulated by altering rejoin hotspot sequences (8). However, the impact of mutations in other rejoin point hotspots, the influence of rejoin hotspot mutations on cbDVG kinetics, and whether different compositions of cbDVG populations vary in their ability to trigger antiviral responses or interfere with their standard viruses remain unclear. In this study, we sought to examine these questions by specifically assessing the impact of R2 mutations on cbDVG production.

Here, we first introduced a 10U mutation into the R2 region in a recombinant RSV (R2-10U virus). Consistent with poly-U mutation in R1, we observed a dramatic loss of cbDVGs containing R2 rejoin points during infection. Moreover, this mutation resulted in a distinct delay in the initial detection of trailer cbDVGs compared to a recombinant WT virus. Interestingly, we observed the rapid emergence and accumulation of a genotype variant bearing a 2-ribonucleotide deletion within the R2-10U mutation sequence, yielding an R2-8U virus. We then generated an R2-8U recombinant virus to examine its impact on viral replication and cbDVG dynamics. Compared to the R2-10U virus, the R2-8U genotype was stable, had enhanced replication kinetics, and correspondingly showed enhanced generation and/or propagation kinetics of cbDVGs with rejoin points within R1-R3. Functionally, we found no differences in the IFN responses induced by virus stocks enriched with cbDVGs predominately comprised of R2 or R3 cbDVGs. Overall, these findings identify a critical RSV trailer sequence whose mutation modulates both viral replication and cbDVG dynamics and suggest that cbDVG generation, particularly near the trailer, may be an evolutionary tradeoff for rapid viral replication.

## Materials and Methods

### Cells and Viruses

HEp2, A549, HEK293T, and BSRT7 cells were cultured at 37°C 5% CO_2_ with Dulbecco’s Modified Eagle’s Media (Gibco) containing 10% fetal bovine serum (FBS), 1mM sodium pyruvate (Gibco), 2mM L-glutamate (Gibco) and 50μg/ml gentamicin (Gibco). Baby hamster kidney cells, constitutively expressing the T7 polymerase (BSRT7 cells), were additionally cultured with 500μg/mL geneticin (Gibco). All cells were treated with mycoplasma removal agent before experimentation (MP Biomedicals). A549 cells constitutively expressing the RSV N protein (A549-RSV-N) were generated by lentiviral transduction using the pLVX-EF1α-IRES-Puro (Takara) lentiviral expression vector followed by puromycin selection and single clone propagation. A549-RSV-N cells were cultured in 10% FBS tissue culture media supplemented with 1.5 μg/ml puromycin. All recombinant viruses were generated using the RSV reverse genetic system in BSRT7 cells. Viral amplification was performed in HEp2 cells. Viral titration was performed in HEp2 cells or BSRT7 cells where indicated.

### Plasmids

Mammalian expression vectors for RSV N (NR-36462), P (NR-36463), M2-1 (NR-36464), and L (NR-36461) proteins, and the RSV reverse genetic backbone pSynkRSV-line19F (NR-36460) were obtained from BEI Resources. The RSV minigenome constructs used to assess the impact of mutations on cbDVG generation have been previously described (8). R2-10U and R2-8U mutations were introduced into the pSynkRSV-line19F reverse genetic system backbone by site directed mutagenesis. To clone specific RSV cbDVG species, the BsiWI restriction enzyme sequence present in the RSV trailer sequence was utilized. First, site-directed mutagenesis (Agilent) was used to insert the NcoI restriction enzyme site immediately following the T7 promoter in the RSV minigenome R1 construct (8). cbDVG 1563 was reverse transcribed from the total RNA of the passage 9 (P9) lineage 1 R2-10U virus using Small-Rev. cbDVG 1563 cDNA was further amplified using NcoI-HhRbz-RevSmall, targeting the end of the RSV trailer sequence with a 5’ overhang containing the NcoI restriction site and the hammerhead ribozyme sequence, and BsiWI-2-R containing the BsiWI restriction site (see Table S1). The cbDVG 1563 amplicon and the R1 minigenome constructs were restriction digested with NcoI and BsiWI followed by ligation. Sanger sequencing of cbDVG 1563 cDNA indicated that it contained an R2-8U mutation, which was introduced into the cbDVG 1563 expressing plasmid by site directed mutagenesis (Agilent). cbDVG 1887 was reverse transcribed from the total RNA of the 10 lineage 2 WT virus using BsiWI-R, and PCR amplification used NcoI-HhRbz-RevSmall and BsiWI-2-R primers followed by ligation into the digested vector.

### RSV minigenome system

BSRT7 cells were transfected with the RSV minigenome construct plasmids along with the codon-optimized plasmids encoding RSV: N, P, M2-1 and the L proteins, respectively.

Lipofectamine 2000 and plasmids (3:1 lipofectamine 2000 to DNA ratio) were incubated at room temperature for 20 minutes and cells were incubated with transfection complexes at room temperature for 2 hours shaking at 250rpm. Cells were incubated overnight at 37°C in Opti-MEM. The following morning, the cell media was replaced with antibiotics free tissue culture media containing 2% FBS. Cells were harvested 48 hours post-transfection for RNA extraction. Extracted RNA was treated with Turbo DNase I and RNA was re-extracted prior to cbDVG RT-PCR analysis.

### RSV reverse genetic system

BSRT7 cells were transfected with pSynkRSV-line19F (WT), R2-10U, or R2-8U along with the codon-optimized plasmids encoding RSV N, P, M2-1 and the L proteins, respectively.

Lipofectamine 2000 and plasmids (3:1 lipofectamine 2000 to DNA ratio) were incubated at room temperature for 20 minutes and cells were incubated with transfection complexes at room temperature for 2 hours shaking at 250rpm. Cells were incubated overnight at 37°C in Opti-MEM. The following morning, the cell media was replaced with antibiotics free tissue culture media containing 2% FBS. 3 days post-transfection, cells were split and maintained in 2% FBS tissue culture media until 50% of cells were mKate2 positive. This was repeated for 2 additional passages at which time the recombinant viruses were collected. Viruses recovered from the transfection are referred to as passage 0 (P0). The total time from transfection to the collection of P0 viruses was 12 days.

### RSV serial passaging

HEp2 cells were infected with 600uL of P0 for 5 days to generate P1. HEp2 cells were infected with P1 virus at MOI 0.1 for 4 days to generate P2. cbDVG enrichment began at P3. Virus lineages were generated by infecting separate sets of HEp2 cells at MOI 5 with P2 viruses or by rescuing identical viruses from independent instances of the RSV reverse genetic system. Subsequent passages were generated by infecting HEp2 cells at MOI 5 with the preceding passage for each virus lineage for 3 or 7 additional passages. Each MOI 5 passage proceeded for 3 days. cbDVG enrichment was confirmed by RSV cbDVG RT-PCR. To generate virus stocks with low cbDVG content, P2 viruses were passaged 2 or 3 times in HEp2 cells at MOI 0.001 for 5 days per passage to generate the P4 and P5 LD viruses, respectively. RSV cbDVG depletion was confirmed by cbDVG RT-PCR. Virus titer was calculated by determining the fluorescent forming units (FFU)/mL. Briefly, HEp2 cells were infected with serially diluted viral supernatants and fluorescent cells were quantified 24 hours post-infection. Cellular fractions underwent three cycles of quick freezing and thawing prior to viral FFU/mL determination.

### Identification of cbDVGs by RNA deep sequencing and ViReMa/VODKA

RNA was extracted from designated viral stocks via Trizol-LS according to the manufacturer’s instruction. Quality control assessments of RNA samples were performed with the Agilent Bioanalyzer. Library construction proceeded with 200ng of total RNA using the TruSeq Stranded Total Sample Preparation kit (Illumina) with Ribo-Zero ribosomal RNA depletion. WT and R2-10U virus sequencing for Fig. 3 and 5 was performed on a NovaSeq6000 sequencer (Illumina) using a Novaseq SP-100 flow cell (Illumina) generating 100bp single-end reads giving ∼43 million reads per sample. WT, R2-10U, and R2-8U virus sequencing for Fig. 6 and 7 was performed on a NovaSeq X Plus sequencer (Illumina) generating paired-end reads giving ∼20 million reads per end. The obtained reads were aligned to the GRCh38 human reference genome using Bowtie 2 (v2.2.9) (37). The unmapped reads were then applied to ViReMa (v0.25) or VODKA to identify junction reads from cbDVGs according to previous publications (8, 38). For VODKA output, reads were additionally assessed using NCBI nucleotide BLAST and junction positions were compared between the two. Any reads with different junction positions between the two algorithms were removed and the sequences of filtered DVG junction reads were then obtained. To determine the number of consecutive As in the mutation region within DVG junction reads, we first selected DVG junction reads with rejoin points positions between nucleotides 15044 and 15098. Using the “stringr” package in R, DVG reads covering the mutation regions were extracted with the sequence pattern GTT[A]*(:::|TATTT), and the number of consecutive As was subsequently counted. The final cbDVG junction reads from the ViReMa output were used to graph the cbDVG break and rejoin point distribution in R. The reference RSV genome used for ViReMa and VODKA was the whole genome plasmid sequence of the RSV reverse genetic system backbone, pSynkRSV-line19F, or that containing the R2-10U or R2-8U mutations. To obtain the viral reads fully aligning to the viral reference genome, unmapped reads were aligned to viral reference genome via Subread (39). The coverage of viral reads to the whole viral genome as well as mutation regions were visualized and calculated using IGV: Integrative Genomics Viewer.

**Figure 1.**
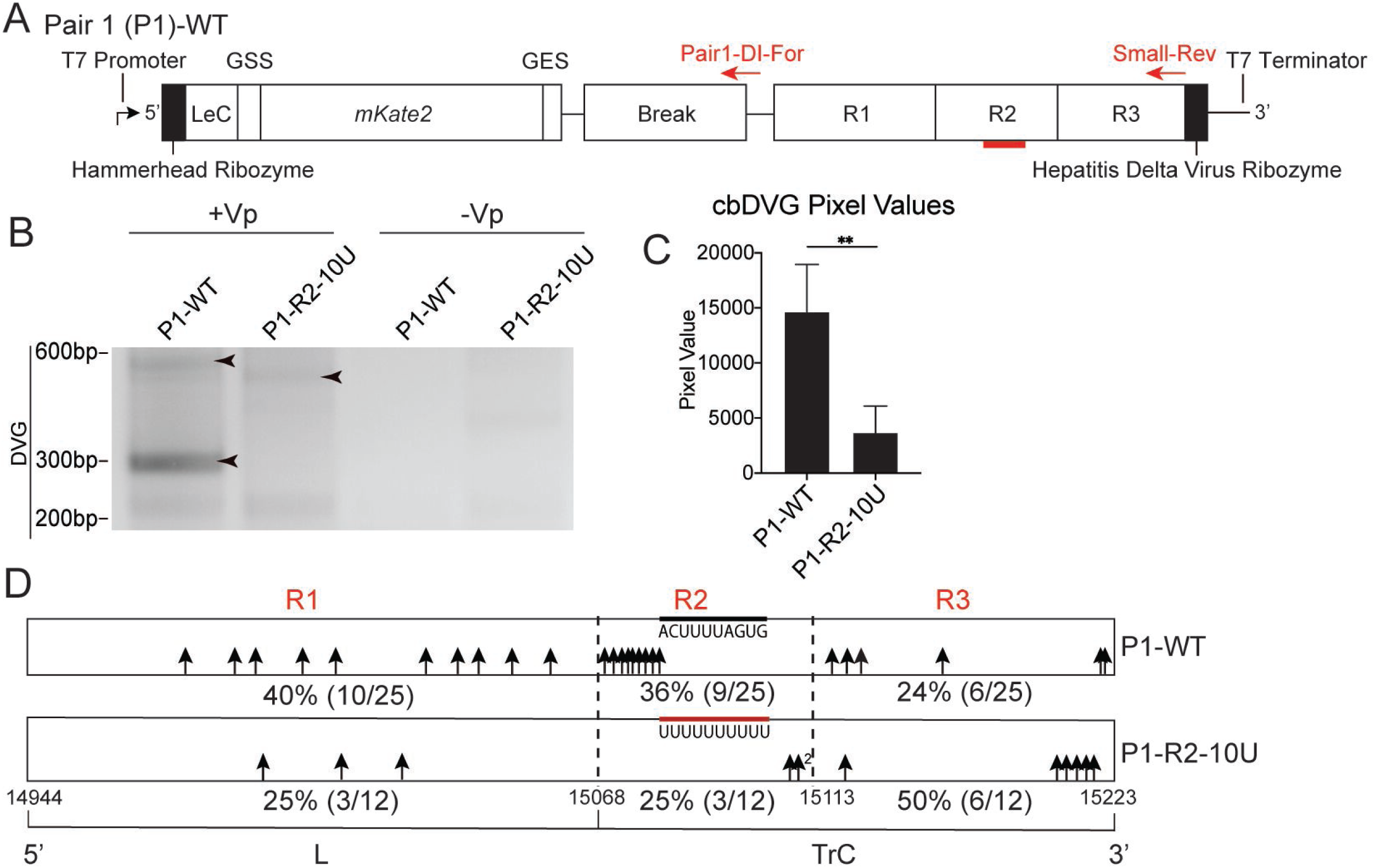
R2-10U mutation attenuated cbDVG near and in the trailer region in an RSV minigenome system. (A) Schematic depiction of the P1-WT minigenome constructs. 5’ to 3’: T7 promoter, hammerhead ribozyme, leader complement sequence (LeC), L gene start signal (GSS), mKate2, L gene end signal (GES), Break, R1, R2, R3, hepatitis delta virus ribozyme, T7 terminator. All constructs are encoded within the pSI1180 vector. The expression of mKate2 first requires synthesis of the negative-sense strand from the trailer to leader mediated by the viral proteins: N, P, M2-1 and L encoded by the four co-transfected helper plasmids. cbDVG generation results from an aberrant round of genomic replication, which can be detected by DVG specific RT-PCR. Bold red lines indicate the location of the 10U mutation within R2. Red arrows indicate the primers, and their locations, used for the cbDVG RT-PCR screen in (B). (B-E) BSRT7 cells were co-transfected with four helper plasmids and P1-WT or P1-R2-10U, respectively, cbDVG RT-PCR using Pair1-DI-For and Small-Rev were performed 48 hpt. (B) Representative agarose gel image of cbDVG-like amplicons detected by cbDVG RT-PCR screening. Black arrowheads indicate cbDVG-like amplicons verified by Sanger sequencing. The sequences of labeled cbDVG-like amplicons are compiled in Figure S2. P1-WT and P1-R2-10U transfections lacking the helper plasmids (-Vp) were included as negative controls. (C) Pixel intensities of cbDVG containing lanes were calculated by subtracting the gel background signal from cbDVG amplicon pixel values for each repeat. **p<0.01 by paired t test, mean±SD, N=5. (D) Schematic summary of rejoin points (black arrows) of all sequence verified cbDVGs from 5 biological repeats. Numbers immediately adjacent to the black arrows indicate the number of cbDVGs with the same rejoin point. Percentage of cbDVGs rejoined in each hotspot is shown beneath the scheme with fractions of the total verified cbDVGs listed in the parentheses.

**Figure 2.**
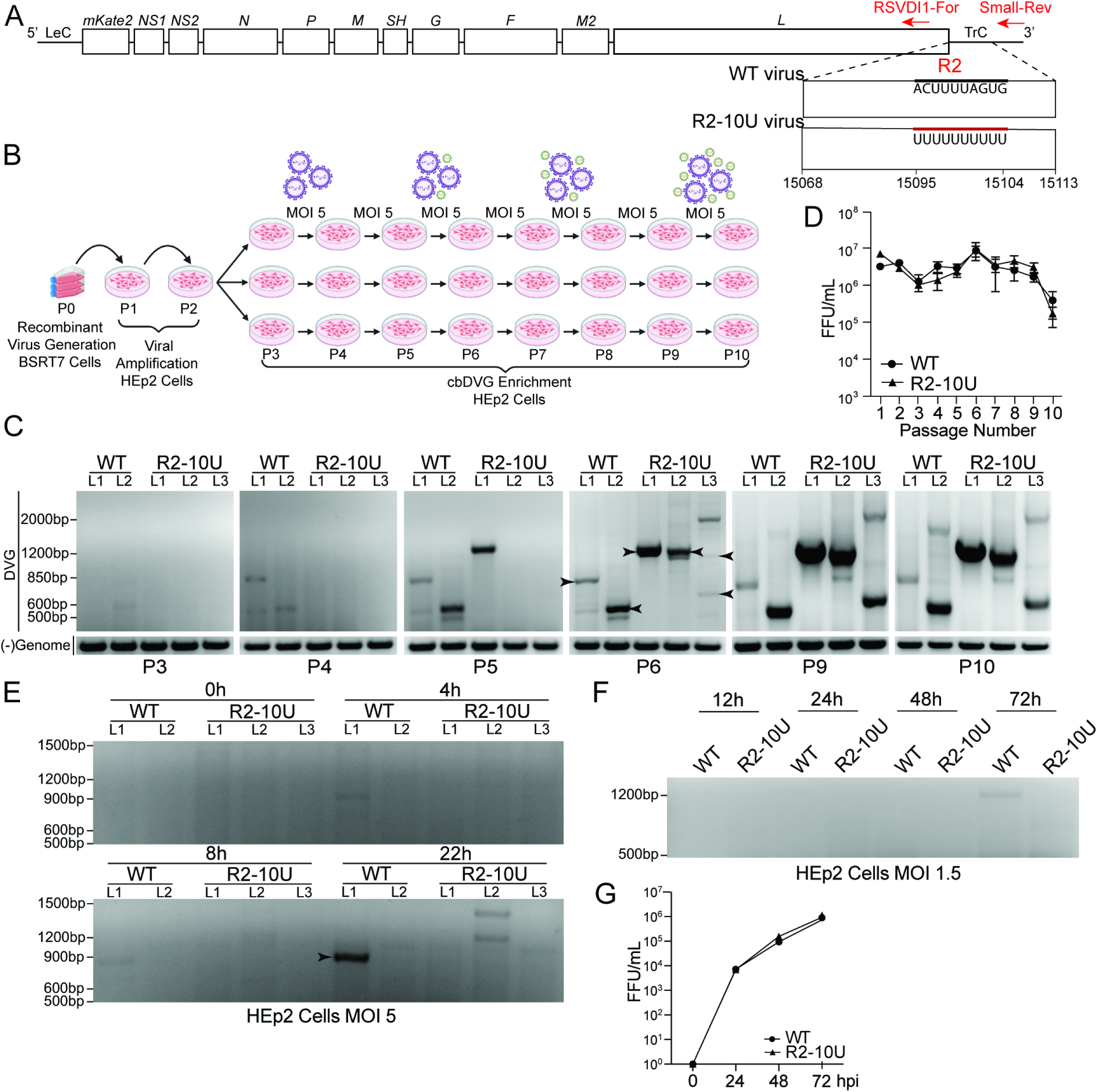
R2-10U mutation resulted in a delayed detection of trailer cbDVGs during infection. (A) Schematic of the recombinant WT and R2-10U viruses. Bold black line indicates the WT sequence and bold red line indicates the R2-10U mutation at antigenomic positions 15095-15104. Red arrows indicate the primers, and their locations, used for cbDVG RT-PCR screening in (C and E-G). (B) Depiction of high MOI cbDVG enrichment strategy. Recombinant viruses were rescued in BSRT7 cells at P0. Recombinant viruses were further amplified in HEp2 cells during P1 and P2. Starting at P3, two lineages of the WT virus and 3 lineages of the R2-10U virus were made, respectively. P3-P10, HEp2 cells were inoculated at MOI 5 for 3 days per passage. Supernatants from each passage were titered and used to generate the subsequent passage. Large purple virus particle icons indicate infectious virus. Small green virus icons indicate defective interfering particles (C) Agarose gel images of cbDVG RT-PCR screens (top) and negative-sense genome RT-PCR (bottom) for P3-P6, P9 and P10 viruses. Lanes corresponding to each lineage for the WT and R2-10U viruses are indicated above. Arrowheads indicate Sanger sequencing confirmed cbDVG amplicons. (D) Virus titers were determined by fluorescent forming units (FFU) for each passage and indicated using FFU/mL. N=2 WT lineages, N=3 R2-10U lineages, mean±SD. (E) Agarose gel image of cbDVG RT-PCR screen of HEp2 cells infected at MOI 5 with WT LD L1-L2 or R2-10U LD virus L1-L3, respectively. Arrowhead indicates Sanger sequencing confirmed cbDVG amplicon with sequences shown in Figure S2. (F-G) Agarose gel image of cbDVG RT-PCR screen (F) of HEp2 cells infected at MOI 1.5 with the WT L2 or R2-10U L2 virus. Virus titers were determined at 24-, 48-, and 72-hpi from the same infection. Viral titration was performed in HEp2 cells (G).

**Figure 3.**
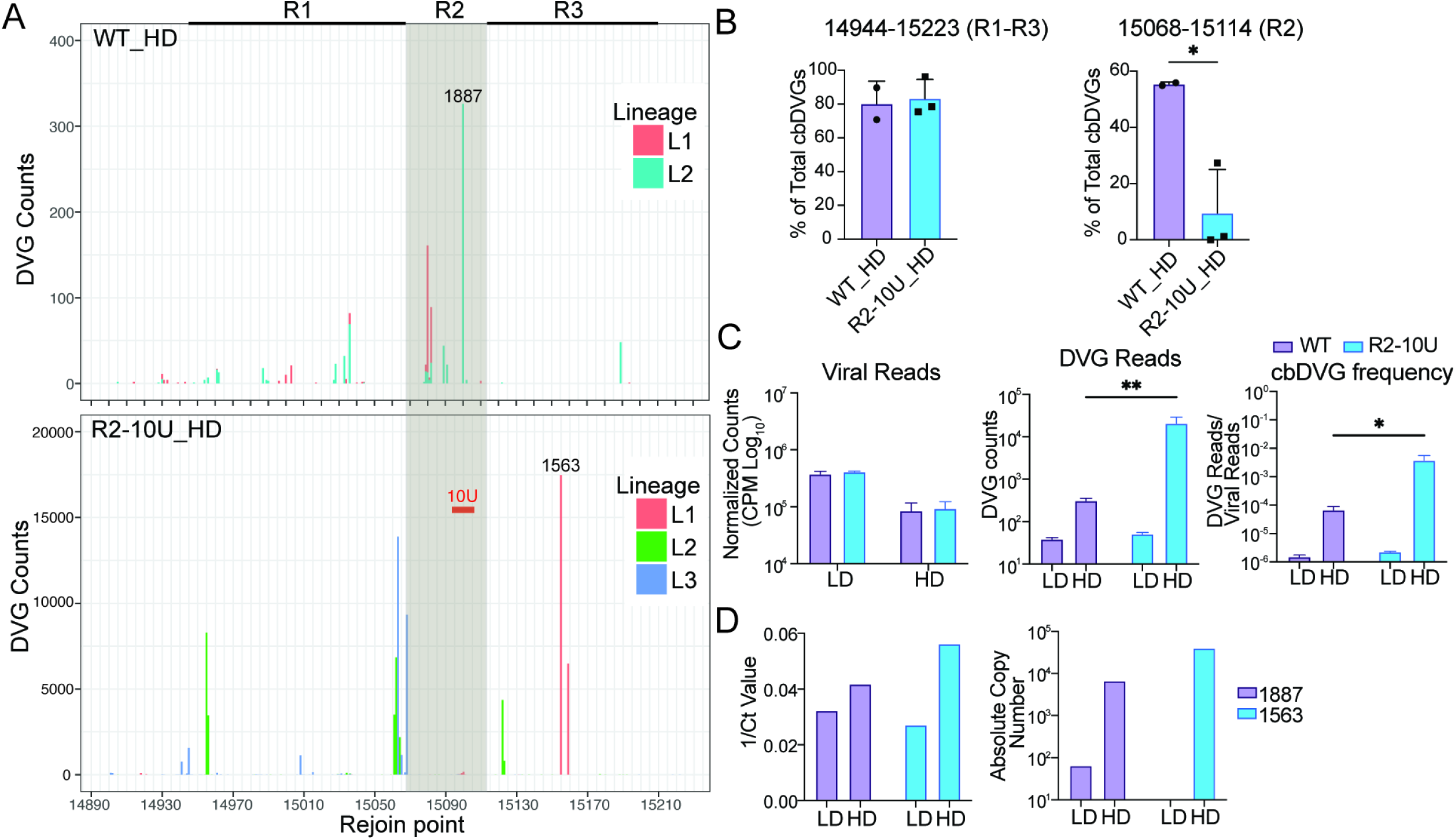
R2-10U mutation diminished cbDVGs with rejoin points at R2 during viral infection. Bulk RNA-seq was performed analyzing the WT and R2-10U P10 HD viruses. (A) Histograms depicting cbDVG rejoin point distributions of the WT and R2-10U viruses. The total number of P10 cbDVG rejoin points at each genomic position spanning 14890-15223 is illustrated. The shaded area indicates the location of the R2 region. The bold red bar within the shaded region indicates the location of the R2-10U mutation. The bolded lines to the left and right of R2 indicate R1 and R3, respectively. cbDVG 1887 and 1563 were selected from the WT and R2-10U HD viruses, respectively, as the dominant cbDVG species for further verification and their rejoin points are indicated on the rejoin point distribution histogram. (B) The percentage of trailer cbDVGs (left) or cbDVGs with rejoin points only at R2 (right) relative to the total cbDVGs were plotted for WT HD and R2-10U HD. *p<0.05 by unpaired t test, mean±SD. (C) Viral reads fully aligning to the RSV reverse genetic system backbone reference genome for the P10 HD and P5 LD of the WT and R2-10U viruses. The total viral reads (left) were normalized to the total sequencing reads shown as log_10_(CPM). Total abundance of cbDVG reads (middle) and cbDVG frequency (right) for the P10 HD and P5 LD WT and R2-10U viruses. cbDVG frequency was calculated by normalizing cbDVG reads to viral reads for each respective lineage of the WT and R2-10U viruses. (D) RT-qPCR validation of cbDVG 1887 and 1563 abundances within WT L2 and R2-10U L3 HD and LD virus supernatants, respectively.

**Figure 4:**
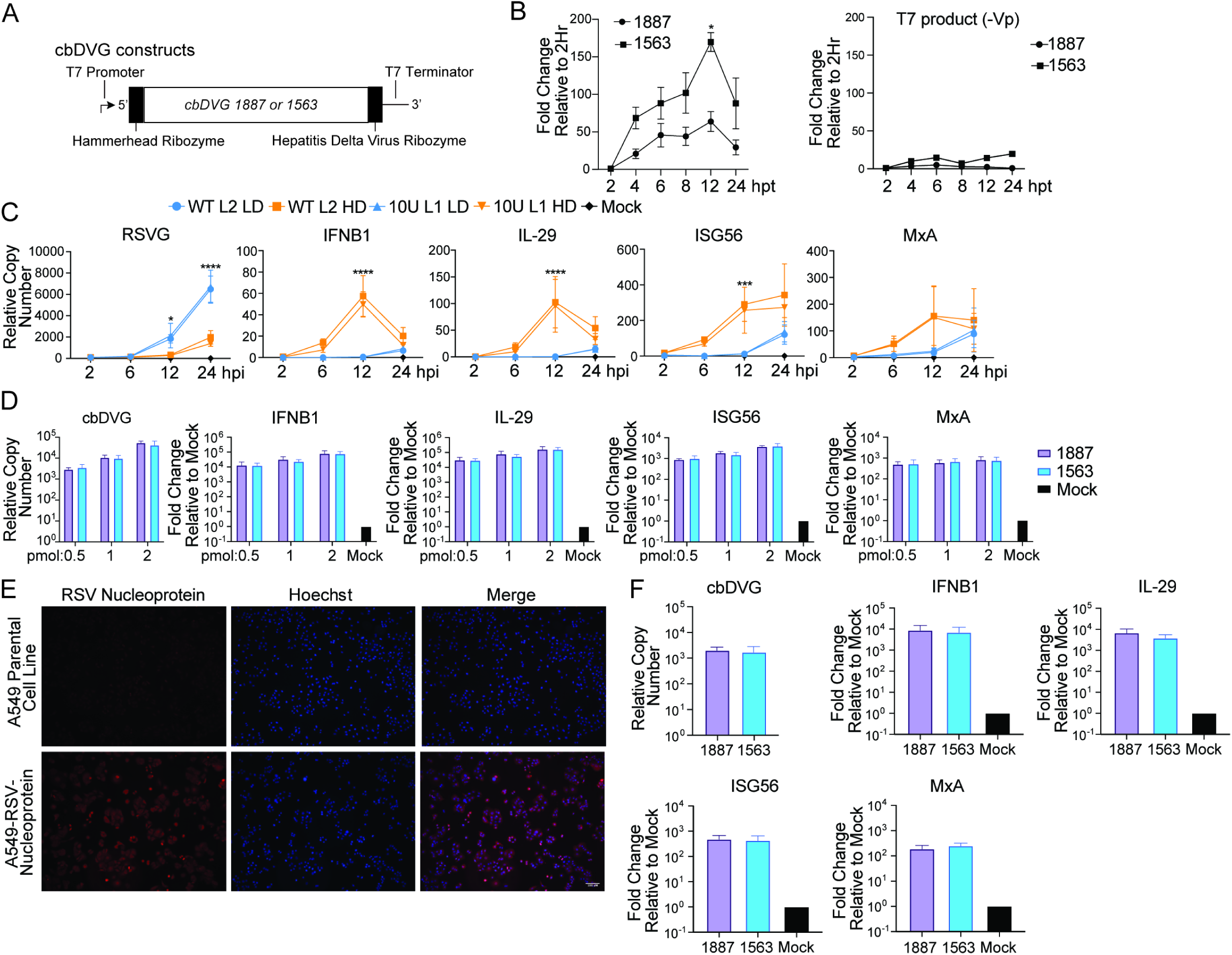
cbDVG 1563 has enhanced replication kinetics and similar immunostimulatory activity compared to cbDVG 1887. (A) Schematic depiction of select dominant cbDVGs 1887 and 1563 expressing constructs. (B, left) BSRT7 cells were co-transfected with four helper plasmids and plasmids expressing cbDVGs 1887 or 1563, respectively, followed by primer specific qRT-PCR (strategy illustrated in S3). cbDVG copy numbers were calculated relative to hamster beta actin and background T7 product values were subtracted. Relative copy numbers were normalized to 2-hours post transfection. *p<0.05 by two-way ANOVA with Sidak’s multiple comparisons test. N=3 independent repeats, mean±SD. (B, right) BSRT7 cells were transfected with plasmids expressing cbDVGs 1887 or 1563 and cells were collected at the same time points, followed by qRT-PCR targeting positive sense DVGs (T7 products). Relative copy numbers were normalized to 2-hours post transfection. (C) A549 cells were infected at MOI 1 with WT L2 HD, R2-10U L1 HD, WT L2 LD or R2-10U L1 LD. The expression of IFNB1, IL-29, ISG56, MxA, and RSV G was examined via qPCR at 2-, 6-, 12- and 24-hpi. Copy numbers were calculated relative to GAPDH. *p<0.05, ***p<0.001, ****p<0.0001 by mixed-effects analysis with Turkey’s multiple comparisons test for RSV G, IFNB1, IL-29, ISG56 and MxA. N=3 independent repeats, mean±SD. (D) cbDVGs 1563 and 1887 were *in vitro* transcribed and 0.5, 1 or 2 pmol of cbDVG RNA was transfected into A549 cells for 6-hours followed by RNA extraction. qRT-PCR targeting IFNB1, IL-29, ISG56, MxA, and cbDVGs 1563 or 1887 was performed. Copy numbers were calculated relative to GAPDH and relative copy numbers were then normalized to mock transfected cells. N=4 independent repeats, mean±SD. (E) A549 cells were transduced to constitutively express the RSV nucleoprotein followed by single clone propagation. Immunofluorescence images using an anti-RSV nucleoprotein antibody for parental and A549-RSV-N cells is shown. (F) A549-RSV-N cells were transfected with 0.5pmol of cbDVG 1563 or 1887 for 6-hours followed by RNA extraction. qRT-PCR targeting IFNB1, IL-29, ISG56, MxA and cbDVG 1563 or 1887 was performed. Copy numbers were calculated relative to GAPDH and relative copy numbers of each gene were normalized to mock transfected cells. N=3 independent repeats, mean±SD.

**Figure 5:**
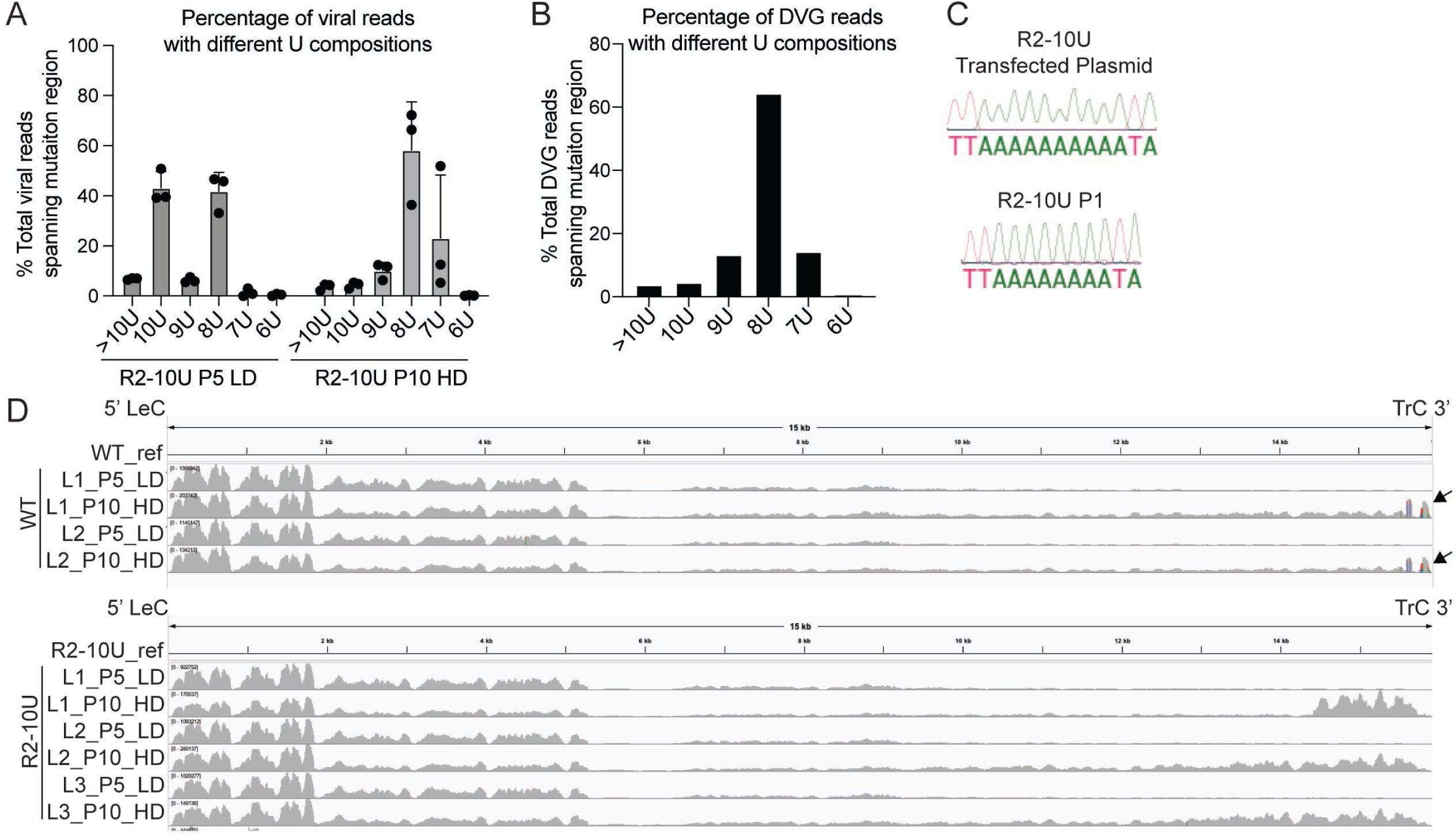
A 2-ribonucleotide deletion emerged in R2-10U virus. (A-B) Among the viral reads (A) or DVG junction reads (B) that aligned to the R2-10U mutation region, the percentage of reads containing 6, 7, 8, 9, 10 or >10 Us were calculated for the R2-10U viruses. (C) Sanger sequencing identified the R2-8U mutation in the P1 R2-10U virus infected cells (bottom) compared to the R2-10U mutation within the plasmid used to generate the recombinant virus via the RSV reverse genetic system (top). (D) RNA-seq reads from the WT and R2-10U viruses were aligned to the WT or R2-10U reference genome, respectively, via subread. The coverages of viral reads were graphed via IGV. Mutations accounting for more than 20% of total reads aligned to that position are colored. Arrows indicate the locations bearing the most variation in WT HD viruses compared to WT reference genome.

**Figure 6:**
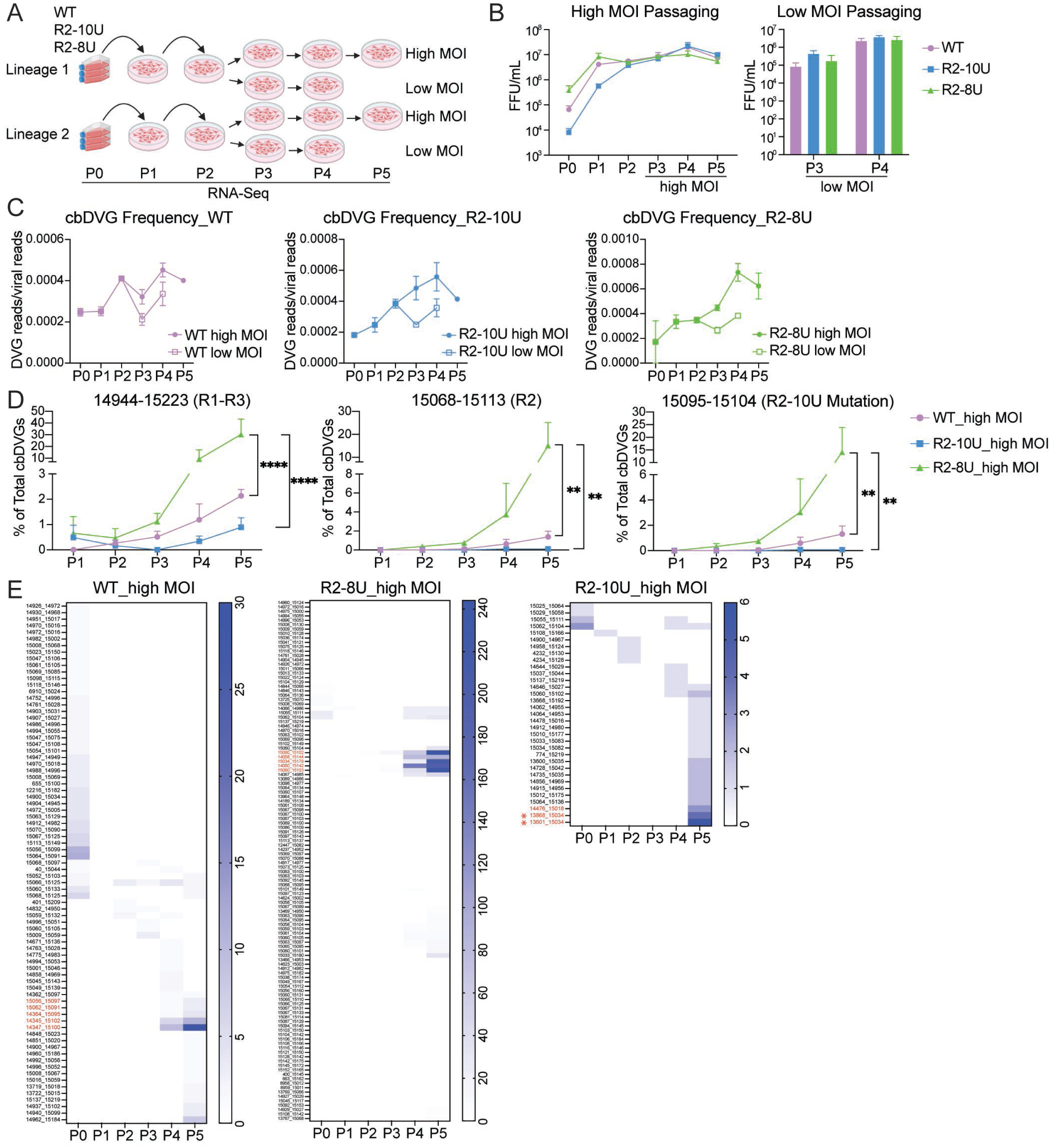
R2-10U virus had reduced trailer cbDVG generation, whereas R2-8U variant enhanced their dynamics compared to WT. (A) Schematic depiction of the strategy used for high or low MOI passaging of WT, R2-10U, and R2-8U viruses. P0 Recombinant viruses, derived from two independent reverse genetic system transfections per virus, were rescued in BSRT7 cells. Recombinant viruses were further amplified in HEp2 cells during P1 and P2. Starting at P3, viruses were passaged either using an MOI of 5 or 0.001. Supernatants from each passage were titered and RNA sequenced. (B) Virus titers were determined by fluorescent forming units (FFU) for each passage either under an MOI of 5 (left) or 0.001 (right). (C) cbDVG frequencies were calculated by normalizing the total cbDVGs reads to the total viral reads for each passages and virus under high MOI or low MOI passaging conditions. (D) Percentages of: trailer cbDVGs (left), cbDVGs with rejoin points at R2 (middle), and cbDVGs with rejoin points in the mutation region (right) relative to the total cbDVG population in WT, R2-10U, and R2-8U viruses across passages. **p<0.01, ****P<0.0001 by two-way ANOVA with Turkey’s multiple comparisons test. (E) Heatmaps of all trailer cbDVGs identified in each passage from WT, R2-10U, and R2-8U. Color intensity indicates the read abundance of each cbDVG obtained from VODKA. *Indicates the major cbDVGs containing R2-8U in DVG junction reads.

**Figure 7:**
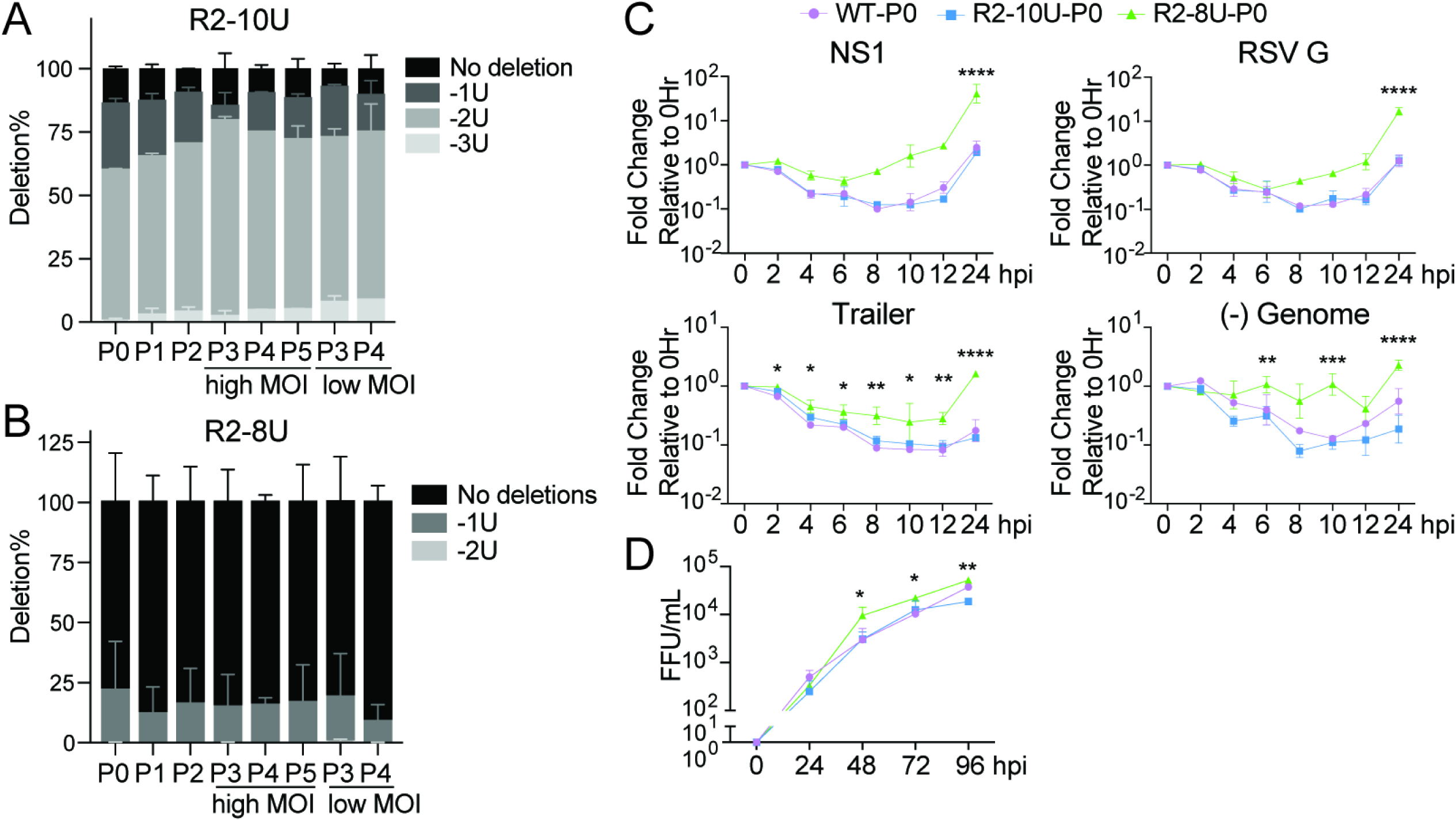
R2-8U has enhanced replication/transcription kinetics compared to R2-10U. (A-B) Among the viral reads from R2-10U (A) or R2-8U (B) that aligned to the R2-10U or R2-8U mutation region, the percentage of reads containing 7, 8, 9, or 10 Us were calculated for each passage. (C) BSRT7 cells were infected with WT, R2-10U or R2-8U P0 viruses at an MOI of 0.0005. The expression of RSV NS1, G, trailer, and (-) sense genome were determined by qPCR at 0-, 2-, 4-, 6-, 8-, 10-, 12- and 24-hpi. Copy numbers were calculated relative to hamster beta actin. Each timepoint was normalized to the 0hpi relative copy number for each gene, respectively. (D) Progeny infectious virus production was determined by FFU/mL quantification at 24-, 48-, 72-, and 96-hpi (bottom left). Viral titration was performed in BSRT7 cells. N=2 biological repeats, mean±SD. *p<0.05, **p<0.01, ***p<0.001, ****p<0.0001 by two-way ANOVA analysis with Dunnett’s multiple comparisons test comparing R2-8U versus R2-10U.

### RNA extraction and RT-(q)PCR for cbDVGs, viral genomes, and host/viral genes

Total RNA was extracted by TRizol or TRizol-LS according to the manufacturer’s specifications. To detect RSV cbDVGs, 1-2 μg of RNA was reverse transcribed using the Super Script III reverse transcription kit (Invitrogen). RNA derived from minigenome experiments was treated with Turbo DNase prior to reverse transcription. The cbDVG RT-PCR screen and primers used has been previously described (8). Briefly, total RNA was reverse transcribed using the Super Script III reverse transcription kit (Invitrogen). Reverse transcription of negative-sense cbDVGs 1563 and 1887 used primer 15292-F. qPCR amplification of both negative-sense cbDVGs used primers 15760-F and Small-Rev. Reverse transcription of positive-sense cbDVGs 1563 and 1887 used primer RSVDI1. qPCR amplification of positive-sense cbDVG 1563 used Small-Rev and 10UL1cbDVG1563-F, while positive-sense detection of cbDVG 1887 used Small-Rev and WTL2cbDVG1887-F. To quantify host/viral gene expression, total RNA was reverse transcribed using the High-Capacity RNA-to-cDNA Kit (Applied Biosystems). cDNA was diluted 1:20-1:40 and amplified with specific primers (Table S1) using the PowerTrack SYBR Green Master Mix (Applied Biosystems). qPCR reactions were performed in triplicate on a ViiA 7 Real-Time PCR System (Applied Biosystems). To quantify the RSV genome, total RNA was reverse transcribed using the Super Script III reverse transcription kit (Invitrogen) using the gRSVDI-F primer. qPCR amplification of the genomic trailer region used gRSVDI-F and Small-Rev. All host/viral genes and cbDVGs were normalized to human GAPDH or hamster beta actin depending on the cell line used. To quantify cbDVG pixel values, pixel densitometry analysis of agarose gel lanes from 100-600bp, with background pixel values subtracted, was performed using ImageJ.

### Immunofluorescence

A549-RSV-N and the parental A549 cell line were seeded on cover slips and cultured overnight. Cells were washed 3X with PBS and fixed with 4% paraformaldehyde for 15 minutes at room temperature followed by 3X PBS washes. Cells were permeabilized in 0.2% Triton X-100 for 10 minutes at room temperature. Cover slips were incubated with 2.5μg/mL Mouse anti-RSV N primary antibody (Abcam ab94806, 1mg/mL) in PBS containing 3% FBS for 2 hours at room temperature. Cover slips were washed 3X with PBS and were incubated with 2μg/mL Donkey anti-mouse Alexa Fluor 594 (Invitrogen A21203, 2mg/mL) in PBS containing 3% FBS for 1 hour at room temperature. Cover slips were washed 3X with PBS and cells were incubated with 1μg/mL Hoechst (Invitrogen H3570, 10mg/mL) in PBS for 7 minutes at room temperature. Cover slips were washed 3X with PBS and mounted on microscope slides using Fluoromount-G (Invitrogen 00-4958-02). Images were taken with a Leica DMIRB inverted fluorescence microscope with a cooled charge-coupled device (Cooke) on Image-Pro Plus software (Media Cybernetics). Images were color overlayed and compiled using ImageJ.

### Functional characterization of RSV cbDVGs

To determine the replication rates of cbDVGs, BSRT7 cells were co-transfected with cbDVG expressing plasmids and plasmids expressing N, P, M2-1 and L. Cells were lysed with TRizol at 2, 4, 6, 8, 12, and 24 hours-post transfection and total RNA was extracted, Turbo DNase treated, and re-extracted. cbDVG specific RT-qPCR was performed and quantified relative to hamster beta actin. To assess the host response to specific cbDVGs, cbDVG 1563 and cbDVG 1887 expressing plasmids were linearized with EcoRI and 1μg of linearized plasmid was used for *in vitro* transcription of cbDVG 1563 and cbDVG 1887 (Agilent RNAMaxx High Yield Transcription Kit) according to the manufacturer’s instructions. A549 or A549-RSV-N cells were transfected with 0.5-2 pmol of RNA for 6 hours at 37°C followed by RNA extraction and examination of cbDVGs, IFN and interferon stimulated gene (ISG) expression by qPCR.

### Statistical analysis

All statistical analyses were performed using GraphPad Prism version 10.0 (GraphPad Software). A statistically significant difference was defined as a p-value <0.05 (based on specific statistical analyses as indicated in each figure legend).

## Results

### R2-10U mutation altered trailer cbDVG rejoin point composition and attenuated cbDVG generation in an RSV minigenome system

Previously, we found that RSV cbDVG rejoin points clustered at the end of the L gene and within the viral trailer complement sequence (genomic positions 14944-15223) after cbDVG enrichment by high multiplicity of infection (MOI) passaging, while break points were widely spread across the antigenome (8). To further characterize the antigenomic region clustered with cbDVG rejoin points, this region was divided into three hotspots with R1 at the end of the L gene, including the L gene end signal, and R2 and R3 in the trailer complement sequence (Figure 1E). The introduction of a poly-U mutation into the R1 region of a recombinant RSV, mutating several key GC ribonucleotides to Us, reduced cbDVGs detected at R1 (8). Here, we sought to determine whether cbDVGs at R2 are also genetically manipulable by introducing a similar mutation in the R2 sequence. We first tested this using a previously developed RSV minigenome system. Briefly, we created a pair1 (P1)-WT construct, under the control of the T7 promoter, containing the RSV leader complement sequence (LeC), an mKate2 reporter gene, the sequence of one previously identified break hotspot (Break), and the wildtype rejoin hotspots R1-R3 (Figure. 1A). The P1-WT 5’and 3’ genomic termini are flanked by the hammerhead and hepatitis delta virus ribozyme sequences, respectively, to generate precise template termini. The (+) sense P1-WT genome is first produced by the T7 polymerase. When co-transfected with helper plasmids encoding the viral factors required for replication and transcription (N, P, M2-1, L), this RNA serves as the template for (-) sense strand synthesis, during which cbDVG generation predominantly occurs and can subsequently be detected by RT-PCR (8).

To test whether the proportion of cbDVGs at R2 is genetically manipulable, a span of ten ribonucleotides (genomic positions 15095-15104) in R2 containing three total G/Cs was first mutated to ten consecutive Us (R2-10U) in P1-WT (P1-R2-10U). BSRT7 cells were then transfected with helper plasmids and P1-WT or P1-R2-10U, and cells were collected for RNA extraction 48 hours post-transfection (hpt). Using cbDVG specific RT-PCR we consistently observed that cbDVG-like amplicons generated by P1-WT were of greater intensity compared to those generated by P1-R2-10U, none of which were detected in the absence of helper plasmids (Figure 1B, primer locations indicated by red arrows in Figure 1A). This observation was further confirmed by pixel density quantification of cbDVG-like amplicons (Figure 1C). To assess how the R2-10U mutation impacts cbDVG rejoin point selection, cbDVG-like amplicons were cloned and their break and rejoin points were determined by Sanger sequencing. For P1-WT, consistent with previous observations, 40% (10/25) of cbDVGs had rejoin points within R1 while 36% (9/25) and 24% (6/25) had rejoin points within R2 and R3, respectively. In contrast, P1-R2-10U had fewer total identified cbDVGs and reduced percentages of cbDVGs containing R1 or R2 rejoin points compared to P1-WT (Figure 1D). To test the transfection efficiencies of P1-WT and P1-R2-10U plasmids, we co-transfected 4 helper plasmids, T7-GFP, with P1-WT or P1-R2-10U, and observed similar percentage of GFP positive cells between P1-WT and P1-R2-10U (Figure S1B). Moreover, we detected more P1-R2-10U templates driven by T7 polymerase than P1-WT in the absence of helper plasmids via qPCR, indicating that fewer cbDVG-like amplicons from P1-R2-10U was not due to less transfection or RNA templates being made (Figure S1C). Taken together, the lesser intensity of DVG amplicons, along with the reduced number of cbDVGs with rejoin points in R1-R3 (herein referred to as trailer cbDVGs) derived from P1-R2-10U, suggest that R2-10U mutation may generally attenuate the generation and/or propagation of trailer cbDVGs.

### R2-10U mutation resulted in a delayed detection of trailer cbDVGs during infection

To determine how the R2-10U mutation impacts trailer cbDVG production and propagation during viral infection, the R2-10U mutation was introduced into the RSV reverse genetic system (R2-10U virus, Figure 2A) (40). Both R2-10U and WT recombinant viruses were successfully rescued and further propagated in HEp2 cells for two passages. To enrich trailer cbDVGs, both passage 2 (P2) viruses were serially passaged at a fixed MOI of 5 until passage 10 (P10). To generate technical replicates, beginning at P3, two separate sets of HEp2 cells were inoculated with the WT virus while three separate sets of HEp2 cells were inoculated with the R2-10U virus (Figure 2B). Supernatants and infected cells were collected and used to monitor virus titers and cbDVG emergence via fluorescent forming unit (FFU)/mL determination and cbDVG specific RT-PCR, at each passage, respectively. For cbDVG RT-PCR, primers specifically targeting trailer cbDVGs were used (indicated by red arrows in Figure 2A). A cbDVG amplicon generated by the WT L2 virus was first detected at P3, similar with previous polyU mutations in R1 (8), and cbDVG amplicons by the WT L1 virus were first detected by P4. Interestingly, for the R2-10U viruses, cbDVG amplicons were first detected at P5 (L1) and P6 (L2 and L3), denoting a striking 2-3 passage delay in cbDVG detection compared to WT (Figure 2C). No differences in virus titer were observed between the WT and R2-10U viruses between P1 and P10 (Figure 2D). To verify this delay, we generated WT or R2-10U virus stocks containing low amounts of cbDVGs (LD) by serial passaging P2 at an MOI of 0.001 for three additional passages. HEp2 cells were then infected with these LD stocks at an MOI of 5 or 1.5.

While high MOI infection encourages pre-existing cbDVGs that competent for efficient replication to further accumulate, lower MOI infection restricts such accumulation, thereby increasing the likelihood of detecting newly generated cbDVGs. As expected, we detected a verified cbDVG for the WT L1 virus as early as 4 hpi under the high MOI condition, while cbDVG amplicons were not detected until 22 hpi for the R2-10U virus (Figure 2E). Given that minimal progeny RSV particles are released before 24 hpi, viral titer determination was not performed. Correspondingly, during the MOI 1.5 infections, a cbDVG amplicon was detected at 72 hpi during WT virus infection, while no cbDVG bands were detected during R2-10U virus infection at any timepoint assessed (Figure 2F). No differences in virus titers were observed at any examined timepoint between the WT or R2-10U viruses (Figure 2G). Altogether, these results suggest that R2-10U virus reduced the generation and/or propagation of trailer cbDVGs compared to WT virus during infection.

### R2-10U mutation diminished cbDVGs with rejoin points at R2 during viral infection

To determine whether the R2-10U mutation impacts the composition of the trailer cbDVG population, total RNA was extracted from the high cbDVG content (HD) P10 viruses for bulk RNA-sequencing. As a control, total RNA was also extracted from LD stocks of each WT and R2-10U virus lineage (P5, used in Figure 2E-G). To identify cbDVG reads, the ViReMa algorithm was used (38). ViReMa identifies the unique break and rejoin junction sequences specific to cbDVGs that do not fully align to the viral reference genome. To assess how the R2-10U mutation impacted the rejoin point composition of the cbDVG population, the rejoin point distributions of cbDVGs spanning the entire genome (Figure S3A) and specifically those within genomic positions 14890-15223 (Figure 3A), containing R1-R3, were plotted for both the WT and R2-10U P10 HD viruses. Consistent with previous observations (8), trailer cbDVGs dominated the DVG population in the WT and R2-10U viruses following high MOI passaging (Figure S3A). Specifically, R2 (54%) was the major rejoin hotspot for WT HD, followed by R1 (20.1%) and lastly R3 (3.97%). For R2-10U HD, however, the frequency of cbDVGs containing R2 rejoin points was significantly reduced relative to R1 and R3 (Figure 3B), indicating that R2 sequences, particularly those within the mutation region, play a key role influencing the generation and/or propagation of trailer cbDVGs containing R2 rejoin points. Like previous results, cbDVG break point clusters were widely distributed across the antigenome ranging from genomic positions 6,000 to 15,000 and these clusters were not conserved among different lineages of the WT and R2-10U viruses (Figure S3B).

Next, to determine the number of reads fully aligned to the viral reference genome (viral reads), the Subread aligner was used (39). To account for potential differences in sequencing depth, viral reads were normalized to total sequencing reads and shown as counts per million (cpm). As expected, viral reads were reduced for both the WT and R2-10U HD viruses compared to their respective LD viruses, indicating that replication competent trailer cbDVGs were sufficiently enriched to reduce the proportion of their respective standard viruses (Figure 3B).

Interestingly, although the dominant trailer cbDVGs in R2-10U HD were detected at a later passage compared to WT HD (Figure 2C), R2-10U-HD had significantly greater total cbDVG reads at P10. This increase remained when cbDVG reads were normalized to viral reads (cbDVG frequency) (Figure 3C). To validate the difference in cbDVG abundance, we examined the absolute amounts of specific major cbDVG species in WT and R2-10U HD stocks via RT-qPCR. Several criteria were established to select the dominant cbDVG species from the WT and R2-10U HD viruses. First, cbDVGs comprising >15% of their respective lineage must have been confirmed by VODKA (Table 1), another algorithm identifying cbDVGs from deep sequencing datasets (8). Next, the select cbDVGs must represent a similar percentage of their lineage according to both ViReMa and VODKA. Finally, for the convenience of primer design, the select R2-10U and WT cbDVGs must contain similar break positions but rejoin points in different rejoin hotspots. Therefore, WT cbDVG 1887, rejoined in R2, and R2-10U cbDVG 1563, rejoined in R3, were selected and specific primers for cbDVG RT-qPCR were designed (Figure S3C). Consistent with the deep sequencing findings, 1563 exhibited greater 1/Ct and absolute copy number values compared to 1887 in their corresponding HD virus lineages (Figure 3D, standard curves in S3D). Overall, these results demonstrated that the R2-10U mutation largely reduced cbDVGs with rejoin points at R2 in trailer cbDVG enriched virus stocks.

**Table 1.**
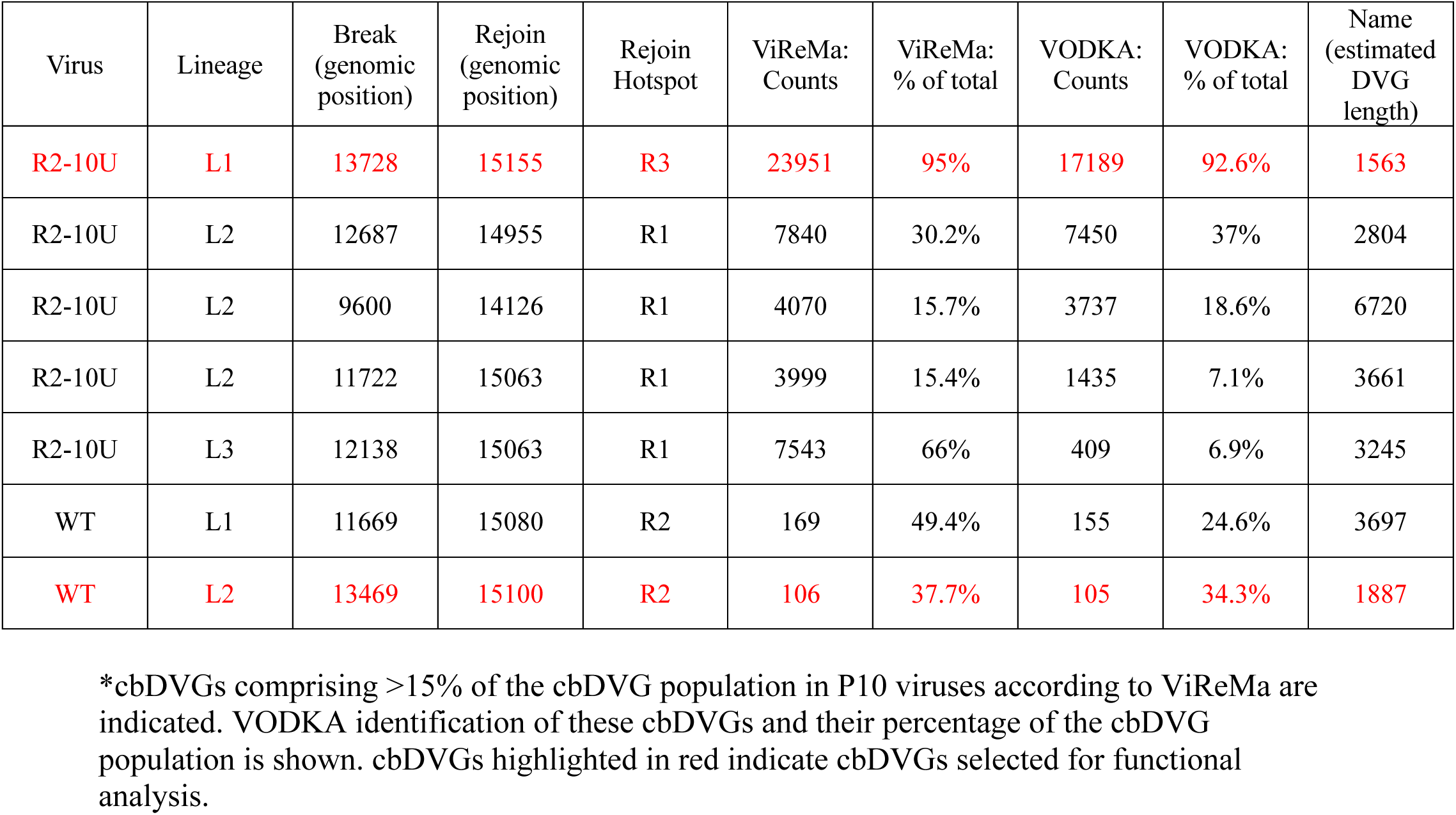
Compilation of major cbDVG species identified by ViReMa and VODKA*.

| Virus | Lineage | Break<br>(genomic<br>position) | Rejoin<br>(genomic<br>position) | Rejoin<br>Hotspot | ViReMa:<br>Counts | ViReMa:<br>% of total | VODKA:<br>Counts | VODKA:<br>% of total | Name<br>(estimated<br>DVG<br>length) |
| --- | --- | --- | --- | --- | --- | --- | --- | --- | --- |
| R2-10U | L1 | 13728 | 15155 | R3 | 23951 | 95% | 17189 | 92.6% | 1563 |
| R2-10U | L2 | 12687 | 14955 | R1 | 7840 | 30.2% | 7450 | 37% | 2804 |
| R2-10U | L2 | 9600 | 14126 | R1 | 4070 | 15.7% | 3737 | 18.6% | 6720 |
| R2-10U | L2 | 11722 | 15063 | R1 | 3999 | 15.4% | 1435 | 7.1% | 3661 |
| R2-10U | L3 | 12138 | 15063 | R1 | 7543 | 66% | 409 | 6.9% | 3245 |
| WT | L1 | 11669 | 15080 | R2 | 169 | 49.4% | 155 | 24.6% | 3697 |
| WT | L2 | 13469 | 15100 | R2 | 106 | 37.7% | 105 | 34.3% | 1887 |

\*cbDVGs comprising >15% of the cbDVG population in P10 viruses according to ViReMa are indicated. VODKA identification of these cbDVGs and their percentage of the cbDVG population is shown. cbDVGs highlighted in red indicate cbDVGs selected for functional analysis.

### R3 cbDVG 1563 has enhanced replication kinetics and similar immunostimulatory activity compared to R2 cbDVG 1887

Our deep-sequencing data indicated that the dominant cbDVGs from R2-10U viruses were enriched to a higher level compared to WT cbDVGs, despite their delayed detection. We hypothesized that this may be due to enhanced replication rates of those dominant cbDVGs enriched in the R2-10U HD viruses. To test this, we compared the replication rates of the two dominant cbDVGs previously selected from the WT (1887) and R2-10U (1563) P10 HD virus stocks. cbDVGs 1887 and 1563 were first cloned under control of the T7 promoter. Specifically, the 5’ and 3’ ends of these cbDVG sequences were flanked by ribozyme sequences to express cbDVGs with accurate termini (Figure 4A). cbDVG expressing plasmids were transfected into BSRT7 cells along with helper plasmids, encoding the viral proteins required for genomic replication, and cells were collected at 2-, 4-, 6-, 8-, 12-, and 24-hpt. To specifically assay for cbDVG replication driven by the viral polymerase, and distinguish from the T7 product, qPCR primers targeting the (-) sense of these cbDVGs were used. By normalizing each timepoint to 2 hpt, 1563 replicated to a higher relative amount than 1887 at each timepoint tested with significant differences between them at 12 hpt (Figure 4B, left). As a control, we also examined the rate of cbDVG templates (+ sense) produced by the T7 polymerase post transfection in the absence of helper plasmids, and observed minimal accumulation (Figure 4B, right), confirming 1563’s greater replication rate compared to 1887.

We next sought to evaluate whether cbDVG populations of differing compositions have distinct IFN stimulation abilities. We first selected WT L2 HD (P10) and R2-10U L1 HD (P10), composed of cbDVGs with rejoin points primarily in R2 (54.4%) and R3 (95%), respectively, for infection in A549 cells at an MOI of 1. A549 cells were alternatively infected with WT LD or R2-10U LD to serve as negative controls. Expectedly, the RSV glycoprotein gene (RSV G) was expressed at higher levels in both LD viruses compared to their respective HD viruses, indicative of cbDVG mediated attenuation of standard virus replication. The expression of IFNB1, IL-29, and ISG56 were significantly elevated at 12 hpi during HD virus infections compared to LD virus infections (Figure 4C). Comparison of WT HD with R2-10U HD, however, did not reveal detectable differences in antiviral gene expression at any timepoint, despite greater cbDVG abundance in the R2-10U virus than WT virus (Figure 4C). To further examine the IFN stimulation of specific dominant cbDVG species identified from the WT and R2-10U HD viruses, we *in vitro* transcribed (IVT) cbDVG 1887 and 1563 RNA and A549 cells were transfected with 0.5, 1 or 2 picomoles. Cells were harvested at 6 hpt followed by RT-qPCR targeting antiviral gene expression and these specific cbDVGs. At each concentration, we detected similar relative copy numbers of 1887 and 1563 indicating similar transfection efficiencies. Relative to mock, at each respective picomolar amount, 1887 and 1563 induced similar expression of IFNB1, IL-29, ISG56 and MxA, indicating that 1887 and 1563 had similar immunostimulatory activities as naked RNAs (Figure 4D). However, cbDVGs are generated during viral replication and therefore should be, at least partially, encapsidated with the RSV nucleoprotein during infection. Consequently, the presence of the nucleoprotein could impact the ability of a cellular RNA sensor to detect cbDVGs. Therefore, we generated an A549 cell line constitutively expressing the RSV nucleoprotein (A549-RSV-N, Figure 4E). A549-RSV-N cells were transfected with 0.5 pmol of IVT cbDVG 1887 or 1563 RNA and antiviral gene expression was assessed. Again, no differences were observed in the expression of IFNB1, IL-29, ISG56 or MxA genes between cbDVG 1887 and 1563 (Figure 4F). Taken together, these data suggest that cbDVGs formed in R2 and R3 likely have similar abilities to induce antiviral signaling.

### A 2-ribonucleotide deletion emerged during R2-10U virus passaging

To assess whether any mutations had arisen in the R2-10U mutation region during virus passaging, reads fully aligning to the R2-10U mutation were compared among lineages of the R2-10U P10-HD and P5-LD viruses. Interestingly, in addition to the original R2-10U sequence, in both P10 and P5 we detected various deletion mutations within the R2-10U sequence including: 1 (9U), 2 (8U), 3 (7U) and 4 (6U) ribonucleotide deletions. We calculated the percentage of reads containing a given deletion relative to the total number of reads aligning to the R2-10U sequence and found that R2-8U, a 2U-deletion variant, already accounted for more than 40% of the population in P5-LD stocks. In P10-HD, R2-8U accounted for roughly 60% of the aligning reads (Figure 5A). Reads aligning to the mutation region could originate from either the full-length viral genome or the homologous regions of cbDVGs (distinct from the junction regions). Therefore, to specially examine the percentage of R2-8U in the cbDVG population, DVG junction reads from P10-HD stocks were obtained via VODKA and filtered for those spanning the mutation region. Consistent with the viral reads pattern, R2-8U constituted more than 60% of the total DVG reads (Figure 5B). These results imply that R2-8U is the major variant in both full-length viral genomes and the DVG population. To assess when the R2-8U mutation may have arisen, we PCR amplified the viral genomic region containing R2-10U from earlier passages and identified R2-8U as early as P1 via sanger sequencing, before the detection of cbDVGs by RT-PCR (Figure 5C).

To examine mutations outside of the R2-10U region, we aligned RNA-seq reads to either the WT or R2-10U viral reference genomes and visualized aligned files with IGV (Integrative Genomics Viewer, Figure 5D) (41). Compared to LD, all HD stocks had additional coverage peaks at the 3’ end of the antigenome, indicating that those reads were from the homologous portions of cbDVGs when compared to the full-length viral genome. A similar observation was previously reported in Sendai virus HD stocks (42). We observed greater variation in WT than the R2-10U viruses due to mutations identified at 3’ antigenomic trailer of the WT HD stocks (arrows in Figure 5A). Because of their specific locations, it is likely that these mutations were in trailer cbDVGs rather than full length viral genomes. Interestingly, each of these mutations were A to G or U to C, consistent with the RNA editing pattern of adenosine deaminase acting on RNA (ADAR) enzymes and have been previously reported in cbDVGs of measles and human metapneumovirus (43, 44). The remaining mutations were considered minor, given that their WT ribonucleotide counterparts comprised more than 50% of the ribonucleotides at those positions (listed in Table S2).

### R2-10U virus had attenuated trailer cbDVG accumulation, whereas the R2-8U variant enhanced cbDVG dynamics compared to WT

Because R2-8U was the most frequent variant in the R2-10U virus, we sought to determine the impact of the R2-8U mutation on viral replication and trailer cbDVG dynamics. To do this, the R2-8U mutation was introduced into the RSV reverse genetic system to generate the recombinant R2-8U virus. For comparison, recombinant WT and R2-10U viruses were generated again in parallel, with each virus produced from two independent transfections of the RSV reverse genetic system. All viruses were then amplified and serially passaged in HEp2 cells at a fixed MOI of 5 (high MOI) or 0.001 (low MOI) similarly as before (Figure 6A).

Consistently, we found no significant differences in virus titers among WT, R2-10U, and R2-8U viruses at any tested passages (Figure 6B). To assess the emergence and accumulation dynamics of trailer cbDVGs following serial passaging in different viruses, we deep sequenced total RNA from each passage, including P0. First, to examine the total amount of cbDVGs, we calculated the cbDVG frequency and found that high MOI passaging, in general, led to greater cbDVG accumulation relative to low MOI passaging (Figure 6C). We then analyzed the distribution of cbDVG break and region point positions across the entire viral genome and observed that cbDVGs of large predicted sequence sizes with rejoin points near the leader end (genomic positions 0-4500) were more abundant than trailer cbDVGs in the early viral passages (Figure S4A). A similar observation was previously reported by Felt et. al (45). Consistent with their report, we observed that trailer cbDVGs only accumulated following high MOI passaging (Figure 6D vs S4B and S4C). Importantly, under high MOI passaging conditions, the rate of proportionate trailer cbDVG accumulation was least in the R2-10U virus, intermediate in the WT virus, and significantly greater in the R2-8U virus. This trend was particularly pronounced for cbDVGs with rejoin points specifically at R2 or within the mutation region (Figure 6D). These results not only validated the delayed detection of trailer cbDVGs during R2-10U virus infection relative to WT by DVG specific RT-PCR (Figures 2 and 3) but also underscored the enhanced dynamics of trailer cbDVG during R2-8U infection. Finally, to determine the passages wherein trailer cbDVGs that later dominate their cbDVG respective populations first emerged, we generated heatmaps of all identified trailer cbDVGs from each virus based on their abundance in each passage. The major trailer cbDVGs in P5-WT, P5-R2-8U, and P5-R2-10U stocks first appeared at P4, P3, and P5, respectively (highlighted in red, Figure 6E). Moreover, the R2-10U viruses displayed the fewest cbDVG species and the lowest abundance per cbDVG, whereas the R2-8U viruses showed the greatest number of cbDVGs and abundance per cbDVG (Figure 6E). Taken together, these data indicated that the R2-10U virus contained few R2 derived cbDVGs and had delayed kinetics of trailer cbDVG accumulation, while its R2-8U variant reversed this phenotype, exhibiting an enhanced ability to produce and/or propagate trailer cbDVGs.

Among the viral reads spanning the mutation region from the R2-10U virus, we again observed the R2-8U genotype as early as P0, consistently accounting for >50% the total viral population across all passages, regardless of high or low MOI passaging conditions (Figure 7A and S5A). In contrast, the R2-8U genotype remained relatively stable in the mutation region, with one replicate maintaining >95% and the other maintaining >75% (∼25% 1U deletion variant) R2-8U (Figure 7B and S5B). Next, we similarly analyzed the DVG junction reads from the R2-10U virus following high MOI passaging. However, due to the limited number of trailer cbDVGs identified in R2-10U through P5, we identified only 7 reliable DVG junction reads spanning the mutation region. Nonetheless, of these reads four contained R2-8U and one contained R2-10U. Notably, both of the two major trailer cbDVGs in R2-10U P5 carried R2-8U (marked with * in Figure 6E). In summary, consistent with our previous observations, these two additional independent repeats strongly support our findings that the R2-8U genotype emerged early during infection and was the major variant in both full-length viral genomes and cbDVGs within the R2-10U population.

We then asked what drove the emergence of R2-8U during R2-10U virus infection and enabled the R2-8U genotype to more rapidly produce and propagate trailer cbDVG. We hypothesized that the R2-10U genotype negatively impacted genomic replication, leading to the emergence and selection of a variant with relatively enhanced genomic replication whose enhanced replication further accelerates cbDVG dynamics. Because R2-10U P0 contained 59.9% R2-8U, the least of any passage, P0 viruses were selected for low MOI growth curves. Additionally, because R2-8U emerged and accumulated in BSRT7 cells, BSRT7 cells were subsequently infected with WT, R2-10U or R2-8U P0 viruses at an MOI of 0.0005. Infected cells were collected at 0-, 2-, 4-, 6-, 8-, 10-, 12- and 24-hpi, followed by RT-qPCR to examine viral replication and transcription levels. Relative to 0 hpi, the WT and R2-10U viruses exhibited comparable levels of RSV NS1, G, and the trailer region. In contrast, R2-8U demonstrated higher expression of these targets at every time point tested (Fig. 7C). qPCR specifically targeting the negative sense viral genome showed a similar trend (Fig. 7C), indicating that the R2-8U virus had enhanced replication/transcription activity compared to WT and R2-10U at the early stages of infection. Virus titers were additionally determined at D1, D2, D3, and D4 post infection and showed that the R2-8U virus had modestly increase compared to R2-10U at 48-hpi, and each timepoint thereafter (Figure 7D). It is worth noting that the average replication rate of the heterogeneous R2-10U virus population was always comparable to that of the WT virus by both qRT-PCR and multicycle growth curve analysis. In summary, these data indicate that R2 region mutations in the RSV trailer sequence impact early viral genomic replication, the kinetics of cbDVG emergence and their subsequent accumulation dynamics.

## Discussion

Historically, cbDVGs were considered a stochastic consequence of viral genomic replication driven by an error-prone RNA dependent RNA-polymerase exclusively during *in vitro* infection (46). However, several reports analyzing the cbDVG distribution profiles from different *mononegavirales* members observed that the rejoin points of enriched cbDVGs clustered at the 3’ end of the antigenome (7, 30, 31), arguing against the completely random generation theory. For RSV, we previously identified conserved cbDVG rejoin hotspots (R1-R3) at the end of the L gene and within the 3’ trailer of the RSV antigenome after higher MOI passaging. Here, we refer to cbDVGs containing a rejoin point within R1-R3 as trailer cbDVGs. We demonstrated that mutating key G/C ribonucleotides at R1 can reduce trailer cbDVGs detected in R1 (8). Importantly, recent work has begun exploring the evolutionary and experimental factors impacting cbDVG dynamics and population composition. For example, Felt et al. observed that cbDVGs of a large predicted sequence length, with break and rejoin points proximal to the 3’ leader end of the genome, predominate in early passages (45). In conditions favoring low levels of viral replication, such as low MOI passaging, these large cbDVGs are maintained but do not accumulate to a significant degree. However, sustained high levels of viral replication, such as high MOI passaging or infection in STAT1 KO cells, promote the diversification of the cbDVG population and emergence and accumulation of trailer cbDVGs (45). Consistent with this report, we observed that these large cbDVGs did not accumulate regardless of MOI and that the accumulation of trailer cbDVGs only occurred in the high MOI passaging condition. Additionally, the enhanced replication kinetics of R2-8U was associated with the faster accumulation of trailer cbDVGs in earlier passages, compared to WT and R2-10U. Overall, our observations support the notion that variables promoting enhanced levels of viral replication promote trailer cbDVG emergence and accumulation.

Recent evidence indicates that not only are DVGs generated in patients, but that DVGs, and their kinetics, also play key roles mediating infection outcomes in patients (23, 25). For example, trailer cbDVGs have been detected by RNA-seq analysis of nasal samples from hospitalized RSV-positive pediatric patients (8). Likewise, recent deep sequencing of brain autopsy samples derived from an individual with subacute sclerosing panencephalitis (SSPE), a progressive neurological disease caused by a persistent measles virus infection, detected numerous cbDVG species with rejoin points clustering at the trailer end of the viral genome (22). While the functions of trailer cbDVGs during human infection remain to be determined, these data highlight the clinical relevance of these cbDVGs in humans. Here, we focused on trailer cbDVGs and showed that cbDVG generation at R2 was largely attenuated by mutating a sequence containing G and C ribonucleotides to 10 consecutive Us (R2-10U) within the R2 hotspot via both DVG specific RT-PCR and unbiased deep sequencing. Furthermore, such attenuation was associated with delayed trailer cbDVG dynamics during R2-10U virus infection compared to WT (Figure 6). Interestingly, we identified a 2-ribonucleotide deletion variant (R2-8U) that emerged as early as P0 within the R2-10U recombinant virus population. By deep sequencing analyses of the recombinant R2-8U virus, we observed that it not only recovered trailer cbDVG generation and propagation kinetics compared to the heterogeneous R2-10U population, but also accelerated cbDVG dynamics compared to WT. This work provides evidence that the kinetics of trailer cbDVG emergence, in addition to their population composition, can be manipulated by mutating key sequences in rejoin hotpots. Although unstable, the recombinant R2-10U virus, and its delayed cbDVG detection kinetics, provides an opportunity to evaluate how delayed trailer cbDVG dynamics impact the host response and infection outcomes *in vivo*. Likewise, R2-8U provides an opportunity to evaluate how accelerated trailer cbDVG dynamics impact host responses and viral pathogenicity.

Because R2-8U emerged before trailer cbDVGs accumulated to a level sufficient for detection and remained the major viral genotype regardless of the content of trailer cbDVGs, we hypothesized that the emergence of the R2-8U variant was due to greater genomic replication efficiency than R2-10U. Although the R2-10U genotype represents a distinct minority in the P0 R2-10U virus population we reasoned that the P0 R2-8U population, comprised of >75% R2-8U, was sufficiently homogeneous to observe potential replicative differences between these two virus populations and compared to WT. Indeed, a low MOI growth curve revealed that the heterogenous R2-10U population demonstrated similar replication kinetics to WT while R2-8U showed significantly enhanced replication kinetics at timepoints within 24 hpi and modest increases in virus titers after 48 hpi compared to R2-10U. The accelerated trailer cbDVG dynamics of R2-8U are likely driven by its enhanced genomic replication.

While it is technically challenging to distinguish *de novo* cbDVG generation from their subsequent accumulation, our data strongly suggest that cbDVG generation at the R2 region was largely attenuated by the R2-10U mutation. For example, because cbDVG accumulation is a function of multiple rounds of the virus life cycle, given that the RSV minigenome system does not produce progeny infectious virus particles it is likely that the cbDVG population observed in minigenome experiments is more representative of the *de novo* cbDVG population rather than enriched populations. Correspondingly, relative to the WT minigenome, we detected fewer cbDVGs containing R2 rejoin points, none within the R2-10U mutation sequence (Figure 1). cbDVG detection in the minigenome system, however, is limited by PCR primer bias and greater sub-cloning efficiency of more intense agarose gel amplicons. Therefore, we further examined the trailer cbDVGs from P0-P5 viruses by deep-sequencing. Consistently, we detected few cbDVGs with rejoin points in R2, or the mutation region, in the heterogenous R2-10U population (Figure 6). Furthermore, the 2 major trailer cbDVGs (rejoin points in R1) detected in R2-10U P5 contained the R2-8U, rather than the R2-10U, mutation. In contrast, the relatively homogenous R2-8U virus had the earliest emergence of cbDVGs with rejoin points in R2 and specifically within the R2 mutation region. All together, these data strongly suggest that the R2-10U mutation attenuates the generation step of R2 cbDVGs whereas R2-8U mutation restores it.

The terminal 36 nucleotides of the RSV trailer complement sequence constitute the antigenomic core promoter sequence, essential for viral genomic replication (47). Deletion mutagenesis experiments have demonstrated that RSV minigenome constructs and recombinant viruses containing trailer sequence lengths greater than the minimal trailer promoter sequence had more efficient promoter activity than the core promoter alone, indicating that a complete trailer complement sequence has inherently greater promoter activity (48, 49). To our knowledge, the R2-10U mutation is the first evidence that the trailer sequence composition of a full-length trailer, outside of the core promoter, impacts RSV genomic replication. More interestingly, the 2U deletion in this sequence significantly enhanced genomic replication. It has been hypothesized that the trailer sequences beyond the core promoter may support viral polymerase recruitment, polymerase transition to a stable elongation mode, or encapsidation (48). The precise replicative defect and augmentation driven by R2-10U and R2-8U, respectively, remain unclear.

It is worth noting that authentic cbDVG species generated during infection are di/triphosphorylated at their 5’ termini and given their complementary ends, cbDVG species theoretically form a stem loop structure with dsRNA ends in the absence of the nucleoprotein, forming the canonical RIG-I ligand (50). Although the cleavage of ribozyme sequences at both termini theoretically yields cbDVGs with accurate terminal ribonucleotides, cleavage results in cbDVGs lacking a 5’ di/triphosphorylated terminus. Additionally, unless the desired IVT products are specifically purified, the ribozyme cleavage products inherently remain. LiCl purification post IVT limits the precipitation of small RNA ribozyme cleavage products, thus it is unlikely that the ribozyme cleavage products comprised a significant proportion of the RNA species derived from our cbDVG IVT reactions (Biorad). Previous reports have implicated the 5’ di/triphosphoryl group as a key component of cbDVG recognition by dsRNA sensors and antiviral signaling (51, 52). Although not directly assayed for, our data indicates cbDVGs 1887 and 1563 strongly induced the antiviral response in the absence of 5’ di/triphosphorylated termini. Nonetheless, these data indicate that, in the context of transfection-based experiments, cbDVGs 1887 and 1563 have similar IFN and ISG induction potentials regardless of the presence or absence of the RSV nucleoprotein, suggesting that their IFN stimulating motif(s) are likely in regions common between 1887 and 1563. More studies are needed to further verify this observation during infection by, for example, packaging specific cbDVGs into virus-like particles and supplementing them during LD virus infection.

The demonstration that RSV cbDVG populations are genetically manipulable provided a new avenue of investigation to assess many questions regarding the mechanisms of cbDVG generation and the impact of distinct cbDVG populations on RSV pathogenesis (8). In this study, we provide additional evidence that cbDVG generation can be genetically manipulated and demonstrate that the kinetics of cbDVG generation can be genetically manipulated as well. The R2-10U and R2-8U viruses provide novel tools to assess how cbDVG dynamics affect host responses. Furthermore, we observed that the R2-8U variant had an enhanced genomic replication with modest increase of virus titers, compared to its R2-10U progenitor and the WT virus, and that it accelerated trailer cbDVG emergence and accumulation kinetics. These observations identify a sequence in the RSV trailer region that, when mutated, critically modulated both viral replication and trailer cbDVG generation and propagation. They further suggest that cbDVG, particularly trailer cbDVG, generation may be an evolutionary tradeoff for more rapid viral replication kinetics. Overall, this work expands the repertoire of RSV genetic tools to alter cbDVG composition and kinetics, providing a unique platform to study RSV genomic replication, cbDVG-driven pathogenesis, and the evolutionary significance of cbDVGs.

## Acknowledgments

We thank Dr. Carolina Lopez for providing the RSV minigenome constructs that were further modified by us for this study, and Dr. Brian Ward for providing the pcDNA3.1-eGFP plasmid to examine minigenome transfection efficiencies.

## Author Contributions

Experimentation and data analysis: J.W.B., X.W.; tissue culture: S.C.; bulk RNA-sequencing: J.W.B.; DVG analysis and graphing for all bulk RNA-seq: J.W.B., G.W., Y.S.; manuscript writing: J.W.B., Y.S.; funding support: T.J.M., Y.S.; supervision: T.J.M., Y.S.

## Data availability

Deep sequencing data from passaging experiments is deposited at GEO (GSE281185).

